# Structural basis for far-red light harvesting in a euglenophyte photosystem II supercomplex

**DOI:** 10.64898/2026.08.20.745976

**Authors:** Rameez Arshad, Hadrien Forêt, David Kopečný, Masami Nakazawa, Farzad Hamdi, Héctor Miranda-Astudillo, Panagiotis L. Kastritis, Pierre Cardol, Roman Kouřil

## Abstract

Photosystem II (PSII) is in eukaryotic phototrophs is generally considered to operate within a more restricted spectral range than photosystem I (PSI), in which long-wavelength chlorophylls are a well-established feature of the peripheral antenna. Whether eukaryotic PSII can acquire comparable far-red-associated properties through lineage-specific antenna diversification has remained unclear. Here we present a 3.09 Å cryo-electron microscopy structure of the C_2_S_2_M_2_L_2_ PSII supercomplex from *Euglena gracilis*, a euglenophyte species harbouring a secondary plastid and unusual light-harvesting system. We show that the euglenophyte-specific antenna protein LhcE9 occupies the position corresponding to canonical Lhcb5, but in a markedly different orientation that creates a distinct interface with the PSII core, particularly with CP43. Combined structural, spectroscopic, mutagenesis and proteomic analyses support LhcE9 as the stably bound PSII antenna subunit most closely associated with the far-red state in the supercomplex. Excitation-energy-transfer calculations further indicate two fast lineage-specific antenna-to-core routes mediated by LhcE9 and PsbX. Together, these findings reveal an unexpected mode of PSII antenna diversification and provide a structural framework for far-red-associated light harvesting in PSII.

## Main

Photosystem II (PSII) is the pigment-protein complex that catalyses water splitting in oxygenic photosynthesis, releasing molecular oxygen and generating electrons that sustain linear electron flow to photosystem I (PSI)^1,2^. In eukaryotic phototrophs, PSII forms supercomplexes in which the dimeric core (C2) complex associates with a variable complement of peripheral light-harvesting proteins, increasing the absorption cross-section and defining the major pathways of excitation energy transfer to the reaction centre^2,3^. Although the PSII core is highly conserved, the organization and composition of its peripheral antenna are remarkably diverse across photosynthetic lineages, reflecting adaptation to different light environments and evolutionary histories^4–7^. Cryo-electron microscopy (EM) studies of PSII supercomplexes from green algae and land plants have established the overall principles of PSII organization, while also showing that lineage-specific changes in antenna composition can substantially remodel supercomplex architecture^8–16^.

PSI and PSII differ in their spectral properties, a feature that contributes to balanced excitation of the two photosystems under varying light conditions^1,17,18^. In eukaryotic phototrophs, long-wavelength chlorophylls have been characterized predominantly in PSI and its peripheral antenna, where they contribute to excitation equilibration and extend light harvesting into the far-red region^17–26^. In cyanobacteria, far-red photosynthesis is achieved through the incorporation of chlorophyll *f* or *d* into modified photosystems, including PSII, enabling absorption beyond 700 nm^17,18^. By contrast, PSII in eukaryotes is generally considered to operate within a more restricted spectral range than PSI^1,17,18^. Whether this view applies uniformly across photosynthetic lineages, however, remains unclear.

*Euglena gracilis* is well suited to address this question. This euglenophyte acquired its plastid by secondary endosymbiosis with a green alga and possesses a photosynthetic apparatus that differs substantially from that of the primary green lineage^27–30^. Its chloroplasts are surrounded by three membranes, its thylakoids lack stacked grana regions, and its light-harvesting system contains lineage-specific features, including polyprotein antenna precursors and an expanded repertoire of LhcbM-related and LhcE proteins^7,28–31^. Recent biochemical and phylogenomic analyses indicated that LhcE proteins form a distinct antenna family in *Euglena*, bind only chlorophyll (chl) *a* together with diadinoxanthin, and contribute to long-wavelength absorption in the photosynthetic apparatus^32^. In parallel, recent PSI structures from *Euglena* revealed an unusual antenna arrangement enriched in LhcE proteins and provided a structural framework for red-shifted light harvesting in PSI^19,20^. Together, these observations suggested that *Euglena* PSII might also employ a non-canonical antenna organization.

Particularly intriguing is LhcE9, which was identified as a component of the PSII supercomplex, but whose precise position remained unresolved^32^. Previous low-resolution single-particle EM analyses showed that *Euglena* PSII can form C_2_S_2_M_2_L_2_-type supercomplex with three classes of associated LHCII trimers, termed strongly (S), moderately (M) and loosely (L) bound, but did not reveal the binding site of LhcE9 or clarify how this antenna component is associated with the PSII supercomplex^32^. Resolving the PSII supercomplex from *Euglena* is therefore important not only for defining its subunit composition and pigment arrangement at high resolution, but also for understanding how PSII antenna architecture was remodelled after secondary endosymbiosis.

Here we present a 3.09 Å cryo-electron microscopy structure of the C_2_S_2_M_2_L_2_ PSII supercomplex from *Euglena gracilis*. The structure shows that the euglenophyte-specific antenna protein LhcE9 occupies the position corresponding to Lhcb5 in green algae and land plants, but is bound in a markedly different orientation, resulting in a distinct interaction pattern with the PSII core. Combined structural, spectroscopic, mutagenesis and proteomic analyses further indicate that LhcE9 is stably associated with PSII and represents the most likely site associated with the long-wavelength state, a property typically associated with PSI and its peripheral antenna in eukaryotes. In addition, the structure reveals other lineage-specific features of *Euglena* PSII, including an extended PsbM architecture and altered pathways of excitation energy transfer from the antenna to the PSII core. Together, these findings provide a structural basis for far-red light harvesting in PSII and reveal an unexpected evolutionary diversification of PSII antenna organization in a secondary plastid lineage.

## Results

### Overall architecture of *Euglena gracilis* PSII supercomplex

The PSII supercomplex was isolated from thylakoid membranes of *Euglena gracilis* by clear-native polyacrylamide gel electrophoresis (CN-PAGE) and analysed by single-particle cryo-EM (Fig. 1). The final reconstruction reached a resolution of 3.09 Å (Extended Data Fig. 1) and revealed a C_2_S_2_M_2_L_2_-type PSII supercomplex comprising the dimeric core complex (C2) and two copies of each S-, M-, and L-LHCII trimers. In its overall organization, the complex resembles the canonical algal PSII supercomplex, indicating that *Euglena* retains the general green-lineage PSII scaffold despite its evolutionary distance and secondary plastid origin.

A major deviation from the canonical organization is found at the monomeric antenna position corresponding to Lhcb5 in green algae and land plants. In *Euglena*, this site is occupied by the euglenophyte-specific antenna protein LhcE9, whereas the position of Lhcb4 remains conserved (Fig. 1c,d). Thus, the *Euglena* PSII supercomplex preserves the overall algal-like arrangement of the peripheral antenna while replacing one of the canonical minor antenna proteins with a lineage-specific component. This observation identifies LhcE9 as the most prominent non-canonical feature of the *Euglena* PSII supercomplex and points to a distinct organization of the antenna-core interface. In addition, fluorescence measurements of the isolated PSII supercomplex revealed an unusual long-wavelength emission not observed in canonical green algal PSII, indicating the presence of low-energy chlorophyll states (Fig. 4) analysed in detail below.

The high-resolution density map allowed modelling of 17 membrane-intrinsic core subunits per monomer, four plastoquinones, 472 light-harvesting pigments and several structural lipids (Extended Data Tables 1 and 2). The pigment composition of the PSII supercomplex was further assessed by (high-performance liquid chromatography) HPLC analysis (Extended Data Fig. 2). The core complex contains the four major PSII subunits (PsbA-D) together with 13 low-molecular-weight membrane-intrinsic subunits (PsbE, F, H-M, T, W, X, Z, and Ycf12 (Psb30)) and is broadly conserved relative to other eukaryotic PSII structures^9–16,33^. By contrast, the extrinsic oxygen-evolving complex subunits PsbO, PsbP and PsbQ were not resolved, although these subunits were detected in the PSII fraction by mass spectrometry (MS) (Extended Data Fig. 3). Thus, they were likely lost during the elution step, which is consistent with their relatively loose association and frequent loss during purification of PSII complexes from plants and algae^11,12,15,16^. In addition to the non-canonical LhcE9 antenna, the structure revealed other lineage-specific features, including an extended C-terminal motif in PsbM and an additional chlorophyll-binding site in PsbX at the interface between the L-trimer and the core complex (see below).

**Fig. 1.**
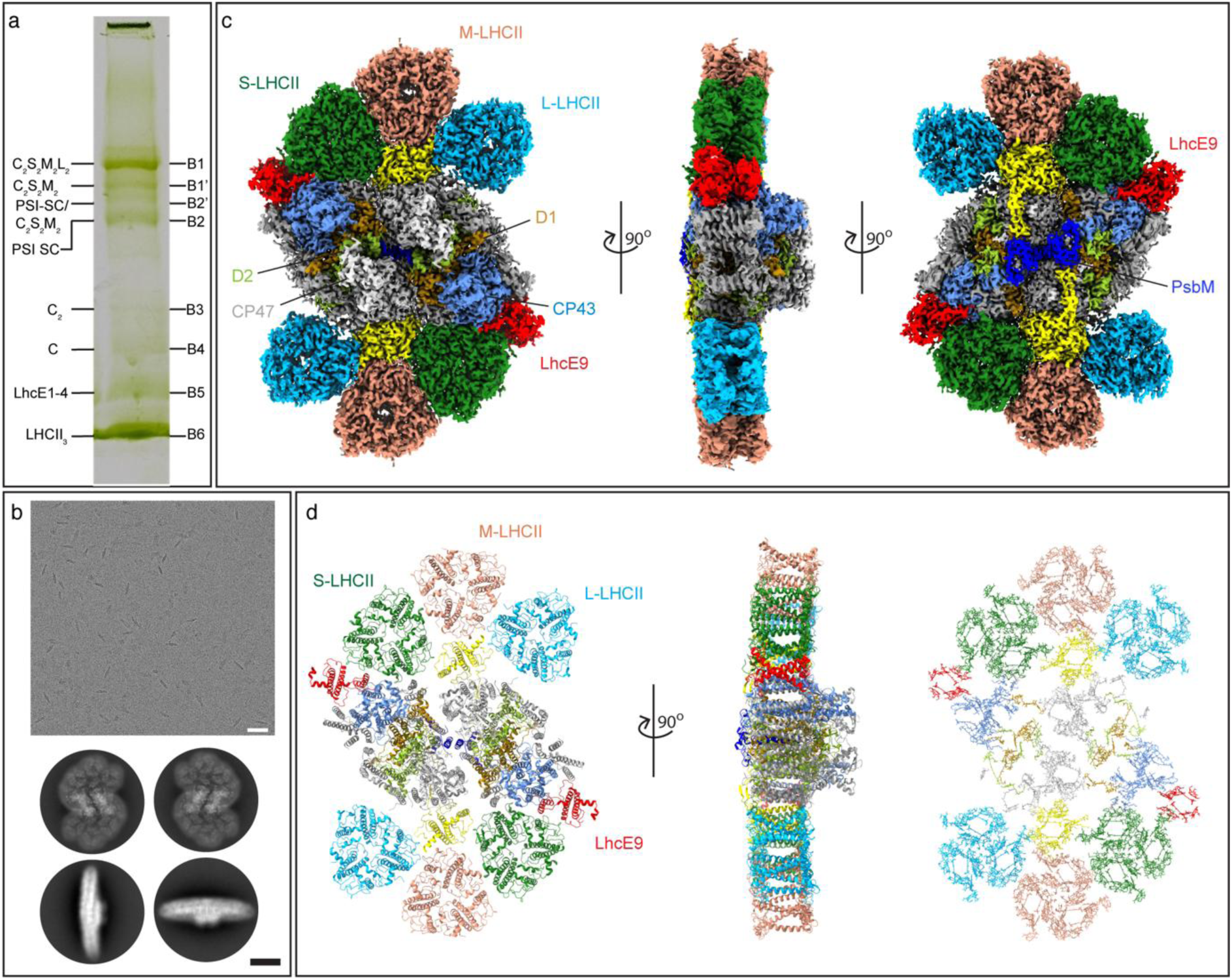
Separation and single-particle cryo-EM analysis of C_2_S_2_M_2_L_2_ photosystem II supercomplex from *E. gracilis.* **a** Clear-native PAGE (CN-PAGE) separation of thylakoid membrane protein complexes isolated from wild-type (WT) *E. gracilis*. The positions of the bands corresponding to PSII-LHCII supercomplexes (B1, B1’), PSI-LHCI supercomplexes (B2, B2’), PSII core complexes (C_2_ and C; B3–B4), free LhcE1-4 antenna complex (B5), and LHCII antenna (B6) are indicated^32^. C_2_S_2_M_2_L_2_ photosystem II supercomplex from *E. gracilis* was obtained from B1 bands. **b** A representative cryo-EM micrograph with specimen from *E. gracilis* PSII supercomplex. The scale bar is 50 nm. The lower section represents the 2D projections of PSII supercomplex from top and side views. The scale bar is 10 nm. **c** Cryo-EM map of PSII-LHCII supercomplex from *E. gracilis*. The large core subunits D1, D2, CP43, CP47 are depicted in orange, green, blue and light gray, respectively. **d** Structural model of *E. gracilis* PSII supercomplex indicating the individual protein subunits and the associated pigments colored according to the coordinating protein chains.

### LhcE9 replaces canonical Lhcb5 and forms a distinct antenna-core module

A characteristic feature of the *Euglena gracilis* PSII supercomplex is the replacement of the canonical minor antenna protein Lhcb5 by the euglenophyte-specific LhcE antenna protein, LhcE9. The LhcE9 protein associates with the core complex in a distinct arrangement involving a translational shift and rotation compared to the canonical Lhcb5 position in plants and algae^8–13,15^ (Figs. 1, 2, and 3). As a consequence, LhcE9 is positioned closer to the PSII core, particularly to the PsbC subunit, thereby reducing the gap between the PSII core and the peripheral antenna (Fig. 2a). Thus, although the overall supercomplex architecture remains broadly algal-like, the canonical Lhcb5 site in *Euglena* is occupied by a structurally and topologically distinct antenna module.

However, the superposition of *E. gracilis* LhcE9 and *C. reinhardtii* (*Cr*) Lhcb5 revealed that LhcE9 is not a simple structural equivalent of Lhcb5. Although the three membrane-spanning helices typical of LHC proteins are retained, LhcE9 appears more compact than canonical Lhcb5, with shorter loop domains and membrane-spanning helices A, B, D, and E (Fig. 2, Extended Data Fig. 4b).

**Fig. 2.**
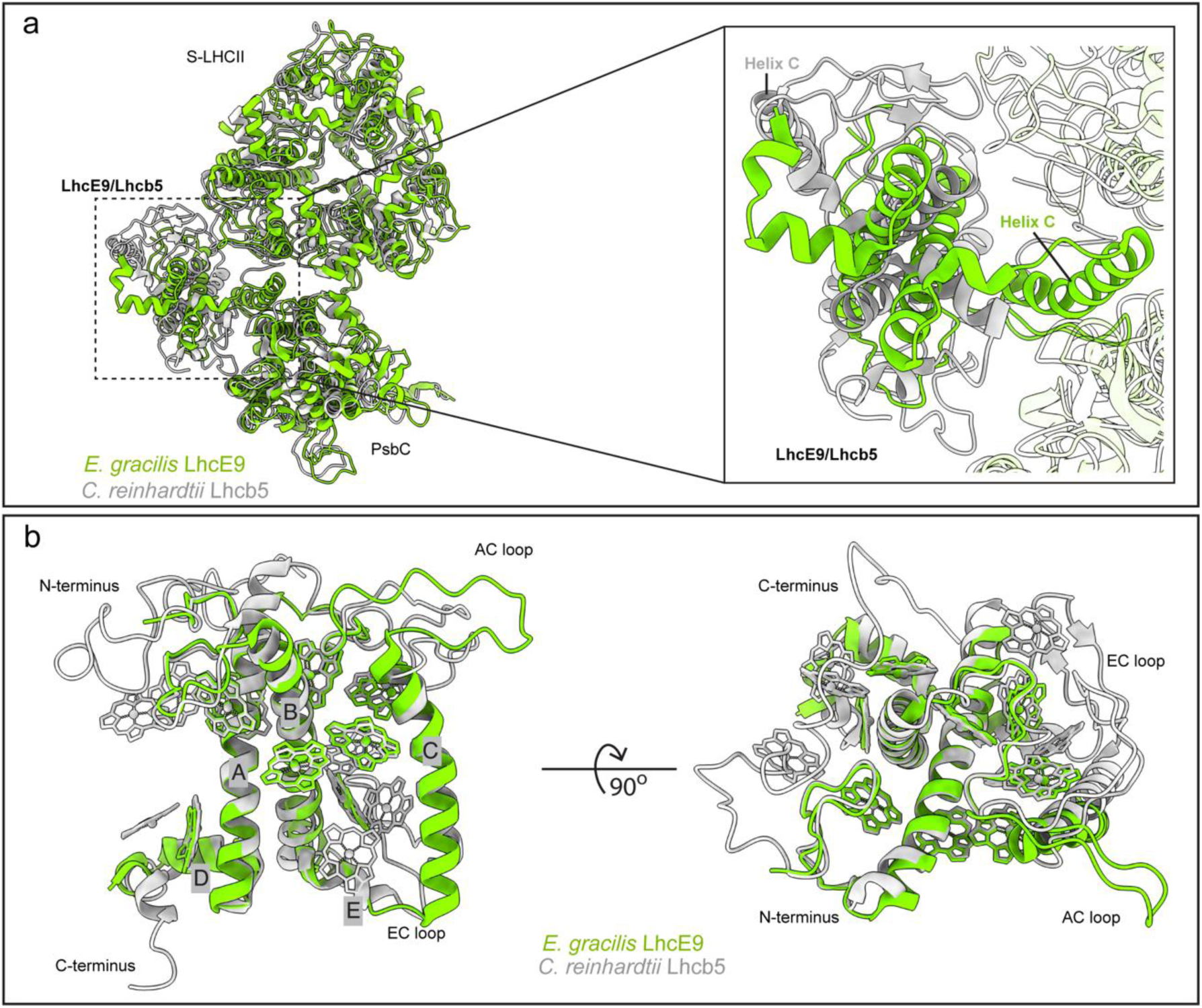
Structural comparison and binding position of LhcE9 and Lhcb5 proteins in *E. gra*cilis and *C. reinhardtii*. **a** Structural alignment of LhcE9 and Lhcb5 binding region involving PsbC and S-LHCII trimer of PSII in *E. gracilis* and *C. reinhardtii*. A large translational shift and rotation of LhcE9 protein brings LhcE9 (through helix C) closer to the PsbC protein. Inset shows a zoomed in view of LhcE9/Lhcb5 binding region of the PSII supercomplexes. **b** Superposition of LhcE9 and Lhcb5 proteins showing relatively conserved structure of transmembrane helices and non-overlapping loop domains.

LhcE9 is also distinct from canonical Lhcb5 in its pigment organization. Based on the cryo-EM density, we modelled eight Chl *a* molecules in LhcE9, consistent with the chlorophyll *a*-only composition characteristic of the LhcE antenna family^32^. Recent PSI structures from *Euglena* showed that LhcE proteins are associated with PSI, indicating that LhcE9 preserves this defining chlorophyll *a*-only feature of the LhcE family despite its integration into PSII^19,20^. Although the positions of the retained pigments remain broadly consistent with the general LHC chlorophyll scaffold, several pigment-binding sites occupied in canonical Lhcb5, including both Chl *a* and Chl *b* sites, are vacant in LhcE9 (Fig. 2b). This reduced Chl *a*-only pigment complement further distinguishes LhcE9 from the canonical PSII minor antenna.

The distinct position of LhcE9 is accompanied by an unusually strong interface with the PSII core. Detailed analysis of the structure showed that LhcE9 interacts predominantly with PsbC/CP43 through helix C and the AC loop, forming a substantial interface of 755.2 Å^2^ (Fig. 3, Extended Data Table 3). On the stromal side, the interaction is reinforced by a network of salt bridges involving Glu481_LhcE9_– Arg153_PsbC_, Glu484_LhcE9_–Arg145_PsbC_, and Glu484_LhcE9_–Arg153_PsbC_, together with multiple hydrogen bonds. On the luminal side, an additional hydrogen bond is formed between transmembrane helix residues Asp450_LhcE9_ and Tyr181_PsbC_. Interestingly, the residues involved in the stromal salt-bridge network are replaced relative to the corresponding positions in *Cr*-Lhcb5 and PSI-associated Lhca proteins (Extended Data Fig. 4a, b), indicating that the LhcE9-PsbC interface is chemically distinct. The predominance of charged residues at this interface generates a locally clustered electrostatic pocket that likely contributes to the stabilization of LhcE9 binding (Fig. 3a). In contrast, the Lhcb5 protein of *C. reinhardtii* interacts with PsbC, PsbZ and the neighbouring LHCII trimer (chains N and Y) with an interface area of 211.5 Å^2^, 337.3 Å^2^, 628.8 Å^2^, and 78.9 Å^2^ respectively . Thus, in *Euglena*, LhcE9 establishes a substantially more extensive and chemically distinct structural interface with the PSII core than canonical Lhcb5. Consistent with structural evidence for enhanced coupling to the PSII core, mass spectrometry analysis reveals that LhcE9 remains stably associated with core complex-containing fractions (Fig. 1, bands 3 and 4), in contrast to CP29/Lhcb4, which shows partial dissociation (Extended Data Fig. 3).

These observations identify LhcE9 as a tightly core-coupled PSII antenna subunit rather than a conventional peripheral linker. The altered docking geometry, the strong electrostatic interface with PsbC/CP43, and the PSI-like aspects of its chlorophyll organization together indicate that *Euglena* has replaced canonical Lhcb5 with a structurally distinct PSII-associated LHC protein that establishes a previously unobserved mode of antenna-core interaction. This unique organization identifies LhcE9 as the most likely structural determinant of the unusual long-wavelength spectral properties of the *Euglena* PSII supercomplex, examined below.

**Fig. 3.**
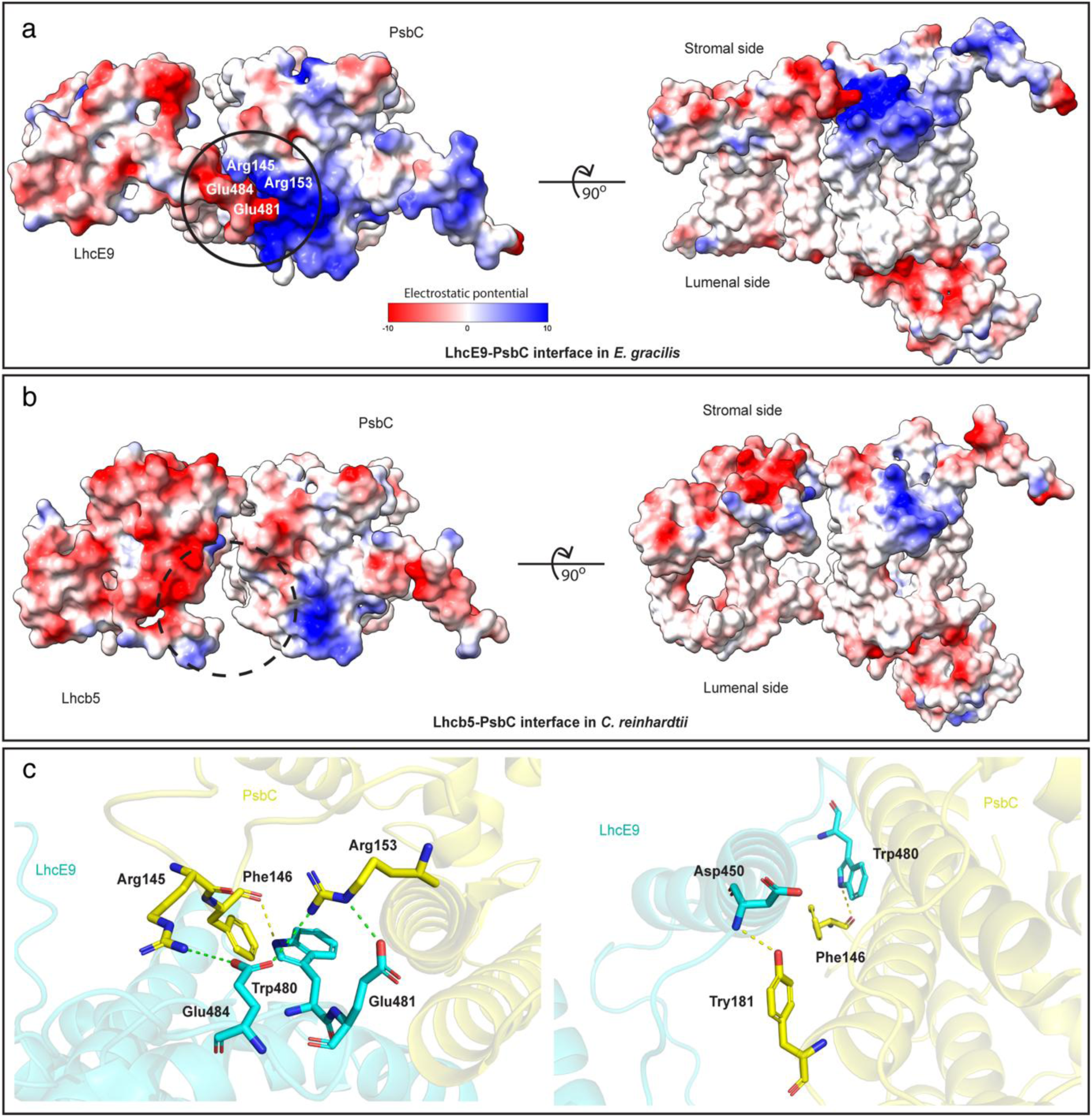
Binding and interactions of LhcE9 and Lhcb5 proteins in *E. gracilis* and *C. reinhardtii*. **a** A representation of electrostatic potential of LhcE9 and PsbC proteins at the LhcE9-PsbC interaction site. The top view from the stromal side and rotated view from the membrane plane show the clustering of the charged (arginine and glutamic acid) residues at LhcE9-PsbC interaction site (black circle). **b** A representation of electrostatic potential of Lhcb5 protein and PsbC subunits. The top and side views show that the Lhcb5-PsbC interaction site is different compared to LhcE9-PsbC proteins with no clustering of charged residues. **c** Interactions of LhcE9 and PsbC protein in *E. gracilis*. The interacting residues are highlighted. Salt bridges are depicted in green, whereas H-bonds are in yellow.

### LhcE9 is associated with the far-red state in the *Euglena* PSII supercomplex

To determine whether this non-canonical antenna organization is linked to the unusual spectral properties of *Euglena* PSII, we combined structural analysis with spectroscopy, CRISPR-Cas9 mutagenesis and proteomics. Fluorescence measurements of the isolated PSII supercomplex revealed a striking long-wavelength emission maximum around 714–717 nm at 77 K, whereas the corresponding PSII supercomplex from *Chlamydomonas reinhardtii* lacked this feature and showed the typical PSII emission maximum in the 685–695 nm region (Extended Data Fig. 5). Room-temperature fluorescence likewise showed a red shift of the *Euglena* PSII complex relative to the *Chlamydomonas* control (Extended Data Fig. 5b). In addition, comparison of room-temperature absorption spectra of the *Euglena* LHCII trimers and PSII supercomplex indicated that enhanced long-wavelength absorption is not a property of *Euglena* LHCII trimers (Extended Data Fig. 6). In combination, these observations indicate that the *Euglena* PSII supercomplex contains additional low-energy chlorophyll states not present in canonical green algal PSII.

A further indication that these long-wavelength states are associated with LhcE9 rather than with canonical LHCII comes from 77K fluorescence analysis of CN-PAGE-resolved thylakoid pigment-protein complexes. In wild-type *Euglena*, the PSII supercomplex band, PSII core-complex-containing fractions, and the free LhcE1-4 antenna complex fraction all exhibited long-wavelength emission around 714–717 nm, whereas the free LHCII fraction showed the shorter-wavelength emission typical of peripheral antenna complexes. By contrast, the PSI supercomplex was shifted further to the red, consistent with the known long-wavelength fluorescence of PSI^19,20,26^ (Fig. 4). Thus, the long-wavelength signal is associated with PSII fractions that contain LhcE9 and is quite similar with the signal associated with free LhcE1-4 antenna complex, supporting the idea that LhcE proteins are the most likely source of the unusual, red-shifted spectral behavior^32^.

To test whether the far-red-associated spectral property of the *Euglena* PSII supercomplex is linked to LhcE9, we analyzed *Euglena* lines lacking CP29/Lhcb4. In these mutants, the PSII supercomplex band was strongly reduced, consistent with destabilization of the peripheral antenna organization (Extended Data Fig. 7). Nevertheless, the long-wavelength fluorescence signal remained clearly detectable in PSII-containing fractions. Thus, the red-shifted spectral property does not depend on the intact outer antenna layer assembled through Lhcb4 and is retained even when PSII supercomplex organization is substantially perturbed in the mutant. Proteomic analysis of the PSII-enriched fractions from the CP29 knockout lines further clarified this result (Extended Data Fig. 8). As expected, Lhcb4 was strongly depleted, whereas the abundance of the major PSII core subunits remained largely unchanged. Importantly, LhcE9 also remained associated with PSII. The persistence of the long-wavelength fluorescence together with the stable retention of LhcE9 argues that the far-red-associated state is more closely linked to LhcE9 than to peripheral LHCII trimers or the minor antenna Lhcb4.

In PSI, long-wavelength or far-red-shifted states have been associated mainly with Lhca proteins in land plants and green algae, where they arise from closely coupled chlorophyll pairs, classically represented by the Chl *a*603/a609 site^17,18,21–26^. In these systems, the *a*603 chlorophyll is typically coordinated by Asn, whereas replacement of Asn by His has often been associated with a blue shift of the far-red maximum^22,23,25^. A similar situation is found in *Euglena* PSI, where a related red-shifted state was assigned to the LhcE6 antenna protein through the Chl *a*305/*a*306 pair, and the corresponding red-shifted chlorophyll (*a*603) is likewise coordinated by Asn^19^. These observations provide a useful reference for interpreting the unusual spectral properties of *Euglena* PSII.

Structural comparison of LhcE9 with red-shifted PSI antenna proteins showed that the putative red chlorophyll pair in LhcE9 is arranged at positions broadly analogous to the long-wavelength chlorophyll pair found in PSI, although the local geometry and protein environment differ (Fig. 5, Extended Data Figs. 9 and 10). In *Euglena* PSII, the analogous chlorophyll in LhcE9 is coordinated by His rather than Asn, yet our spectroscopic data consistently show a red-shifted fluorescence maximum at 717 nm (Fig. 4b, Extended Data Fig. 7b). This indicates that replacement of Asn by His does not necessarily abolish far-red absorption. Similar observations have been reported in other systems. In the moss *Physcomitrium patens*, for example, Chl *a*603 is coordinated by His in Lhca2b, yet the complex still exhibits far-red fluorescence, albeit shifted by about 10 nm relative to land-plant PSI^17,18^. Taken together, these examples indicate that the spectral properties of red chlorophylls are determined not only by the coordinating residue, but also by the geometry of the chlorophyll pair and the surrounding protein environment^18,22–25^.

In this context, structural comparison of LhcE9 with the putative red-shifted LhcE6 site from *Euglena* PSI showed that the candidate red chlorophyll pair in LhcE9 is translationally shifted and rotated relative to the PSI reference^19^. The Mg–Mg distance between Chl *a*804 and Chl *a*807 in LhcE9 is 9.6 Å compared to 8.9 Å between the Chl *a*306/*a*305 pair in the PSI reference (Fig. 5b), a difference that may contribute to weaker excitonic coupling. However, comparison with the *Euglena* LhcbM-containing LHCII trimers indicates that Mg–Mg distance alone is not sufficient to define a far-red site, since several corresponding chlorophyll pairs in these antennae display broadly comparable distances and packing, yet the isolated LHCII fraction emits at 683 nm rather than at 717 nm (Fig. 4b). Thus, the long-wavelength state associated with LhcE9 is unlikely to be explained by Mg–Mg distance alone, but rather by the precise relative orientation and ring-to-ring packing of the two chlorophylls, together with the specific steric and electrostatic properties of the surrounding protein environment. In LhcE9, displacement of helix C, potentially associated with substituted residues and the longer EC loop, is likely to alter the orientation and local packing of the Chl *a*804/*a*807 pair (Fig. 5a). Together, these features provide a plausible structural explanation for why the long-wavelength state in *Euglena* PSII is red-shifted, yet distinct from the more strongly shifted PSI red forms. While the precise far-red-absorbing chlorophylls in *Euglena* PSII cannot yet be assigned unambiguously, the combination of 77 K fluorescence analysis of CN-PAGE-separated fractions, CP29/Lhcb4 knockout analysis, proteomic detection of LhcE9 in PSII-containing fractions and structural comparison supports LhcE9 as the most likely location of the long-wavelength state in the PSII supercomplex.

**Fig. 4.**
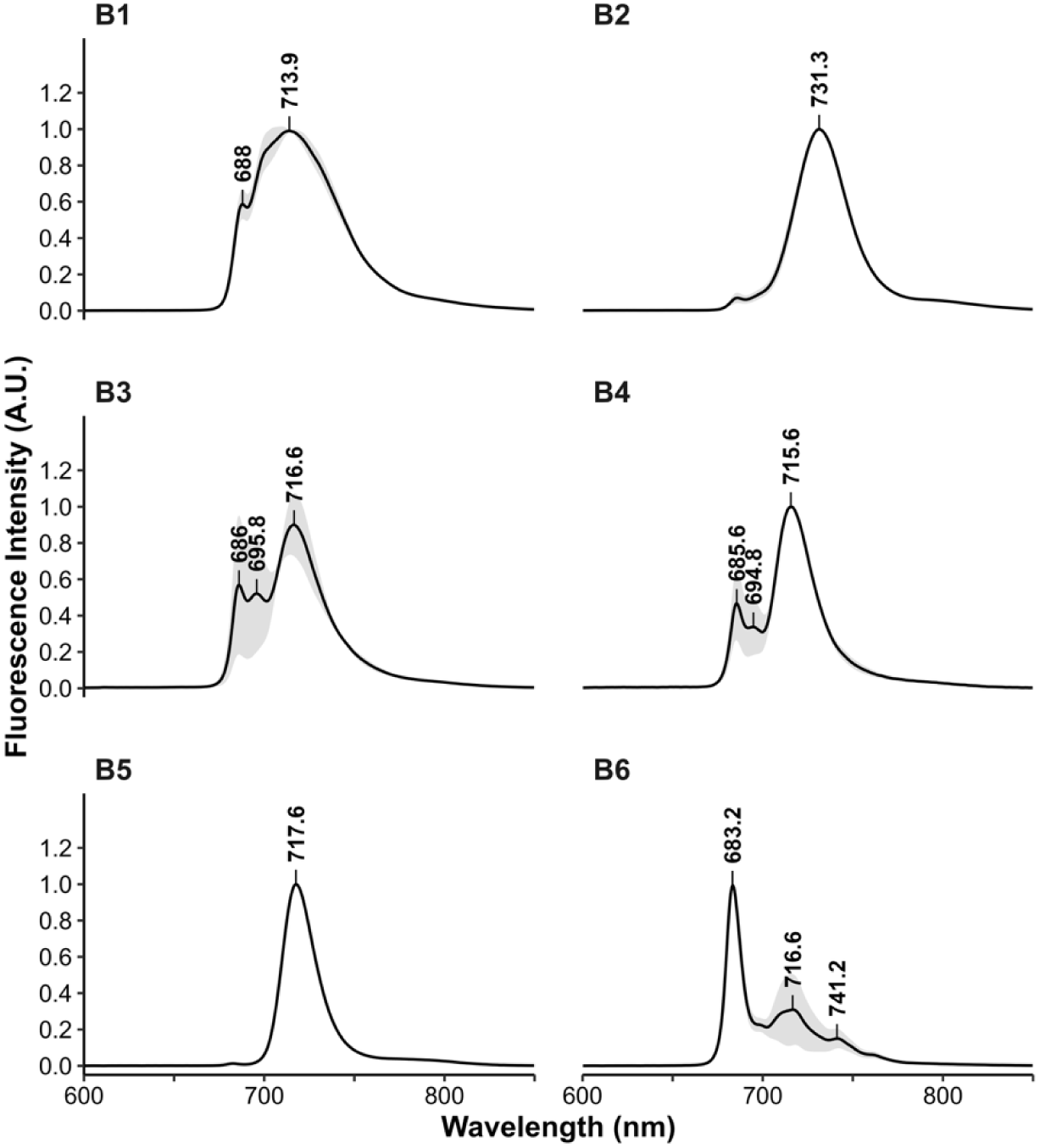
77 K fluorescence spectra of photosynthetic complexes from wild-type *Euglena gracilis*. 77 K fluorescence emission spectra recorded for the individual CN-PAGE bands (B1–B6; see Fig. 1a). The PSII supercomplex band (B1) exhibits emission maxima around ∼714–717 nm, consistent with PSII– antenna assemblies. The PSI supercomplex (B2) shows a red-shifted emission maximum around ∼730– 732 nm, typical of PSI long-wavelength chlorophylls. PSII core complex–containing bands (B3–B4) display emission maxima near ∼717 nm with a shoulder around ∼688–690 nm. The LhcE1-4 antenna complex band (B5) shows long-wavelength emission, supporting the broader association of LhcE proteins with red-shifted fluorescence in *Euglena*. The LHCII band (B6) peaks at ∼683 nm, a wavelength characteristic of peripheral light-harvesting complexes. Fluorescence spectra represent the mean of 3 independent biological replicates. All spectra were normalized at 1 to the maximum.

**Fig. 5.**
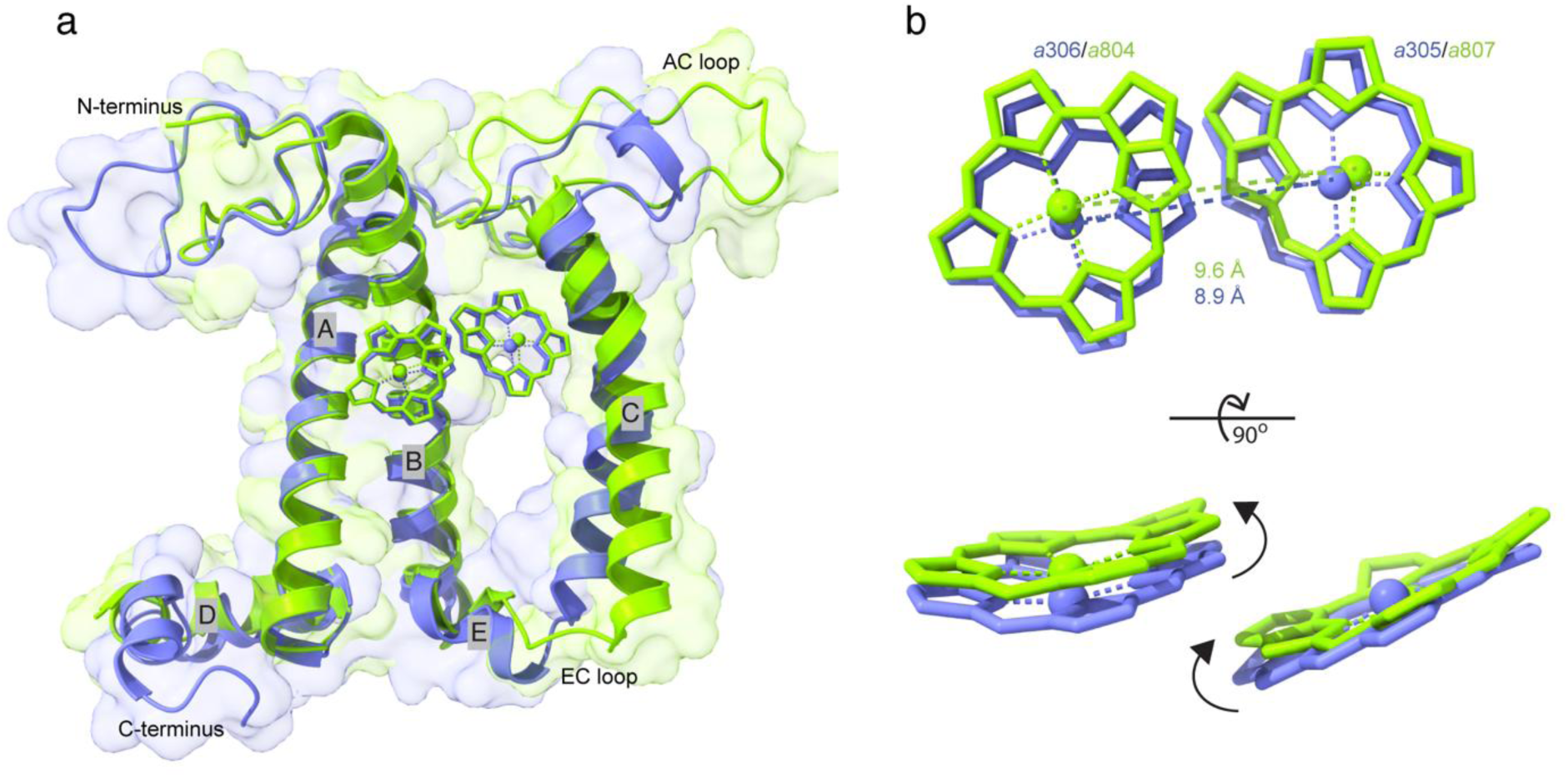
Structural alignment of *E. gracilis* LhcE9 from PSII with LhcE6 from PSI containing red-shifted absorption. **a** Superposition of LhcE9 (green) and LhcE6 protein of PSI (LHC-13, PDB 9VJS/chain j) (blue). The structure of transmembrane helices A and B is conserved, whereas helix C is shifted likely due to longer EC loop. The positions of red chlorophyll pairs in LhcE9 and LhcE6 (9VJS) are shown. **b** An overlay of the positions of red chlorophyll pair in LhcE9 and LhcE6 (9VJS). The distance between Mg–Mg centers and positional difference within red chlorophyll pair are highlighted.

### Excitation energy transfer calculations identify two fast lineage-specific connections to the PSII core

To evaluate the functional consequences of the distinct antenna architecture of the *Euglena gracilis* PSII supercomplex, we calculated excitation energy transfer rates between the individual antenna subunits and the PSII core. Overall, the major pathways of excitation energy transfer are broadly similar to those described for green algal PSII, with energy flowing from the peripheral LHCII trimers towards the inner antenna proteins CP43 and CP47 (Fig. 6)^9–12^. Thus, despite its unusual antenna composition, the *Euglena* complex retains the general functional logic of eukaryotic PSII supercomplexes.

A first distinctive pathway is mediated by the newly identified chlorophyll bound to the PsbX subunit (Extended Data Fig. 11). This chlorophyll lies at the interface between the L-trimer and the PSII core and creates a fast connection between the loosely bound antenna and the core complex. Transfer from L-LHCII chains P and O to PsbX is rapid, and excitation is then transferred predominantly from PsbX to CP47. These calculations identify the PsbX chlorophyll as a potentially efficient bridge linking the peripheral L-trimer directly to the core antenna. Because this chlorophyll site has not been described in canonical green algal or plant PSII structures, it might represent a lineage-specific modification of antenna-to-core energy transfer in *Euglena*^9–14^.

A second distinctive pathway is associated with LhcE9. In contrast to canonical Lhcb5, which is mainly connected to neighboring antenna subunits of the S-LHCII trimer^12^, LhcE9 in *Euglena* is coupled particularly strongly to CP43. Calculated transfer from LhcE9 to CP43 is markedly faster than transfer from LhcE9 to neighboring S-LHCII subunits, suggesting strong antenna-core connectivity rather than as a conventional component of the outer antenna layer. This behavior is fully consistent with the unusual docking geometry of LhcE9, which brings it into close contact with PsbC/CP43 and creates a tight antenna-core interface. Although the calculated transfer rates indicate strong structural connectivity between LhcE9 and CP43, the presence of a low-energy far-red state in LhcE9 may also introduce partial energetic trapping or equilibration within LhcE9, analogous to red chlorophyll states in PSI.

Collectively, these analyses indicate that the excitation-energy network of *Euglena* PSII is broadly conserved relative to green algal PSII but contains two strong lineage-specific antenna-to-core connections: a PsbX-mediated route from the L-trimer to CP47 and a direct LhcE9-CP43 connection. These pathways provide close energetic coupling to both inner antenna proteins of the PSII core. In this context, the position of LhcE9 may be especially significant. If LhcE9 serves as a docking site for an additional LhcE1-4 antenna complex, as suggested previously^32^, its strong coupling to CP43 could provide a route for transferring excitation energy from such an antenna directly to the core. Although this possibility remains speculative, it is consistent with both the structural organization of LhcE9 and the calculated transfer rates.

**Fig. 6.**
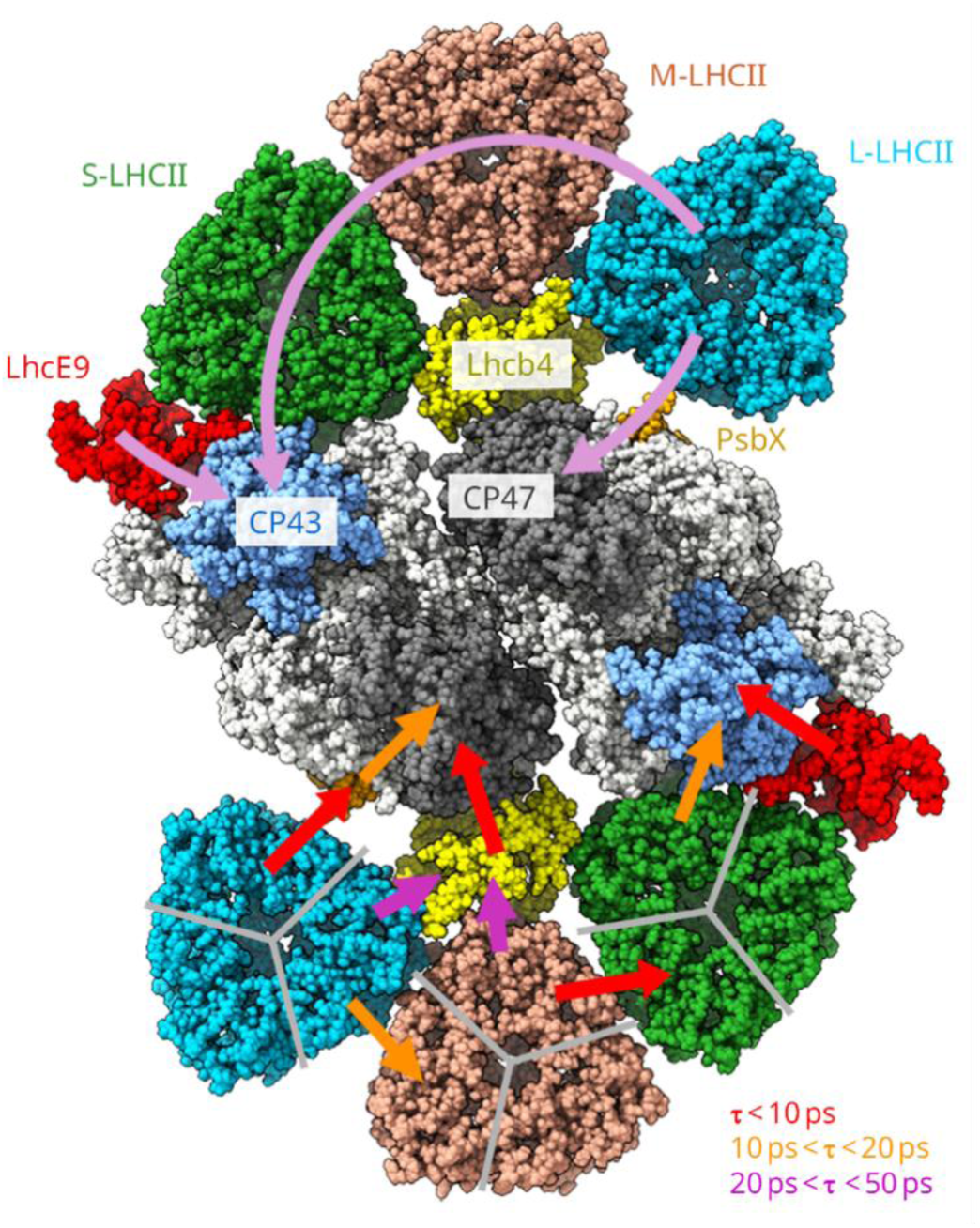
Major energy transfer pathways within the PSII C_2_S_2_M_2_L_2_ supercomplex. The upper part of the PSII model shows the main overall FRET pathway from (i) the LHCII trimers to the PSII core complex (light magenta arc arrow), going from the L-LHCII trimer through the M-LHCII to the S-LHCII trimer and to CP43, the inner antenna of the core complex, (ii) monomeric antenna LhcE9 to CP43, and (iii) L-LHCII trimer to CP47 via PsbX core subunit. At the bottom of the PSII model, the arrows of different colors indicate approximate lifetimes of efficient FRET processes from adjacent subunits of LHCII subunits (L-/M-/S-LHCII trimers, Lhcb4 and LhcE9) towards the core complex (CP43 and CP47). The PSII C_2_S_2_M_2_L_2_ supercomplex is shown from the lumenal side. Source data are provided as a Source Data file.

### Additional lineage-specific features of antenna-core organization in *Euglena* PSII

Although LhcE9 represents the most prominent antenna innovation in the *Euglena gracilis* PSII supercomplex, additional structural features also distinguish this complex from canonical green-lineage PSII. One of the most striking differences is found in the low-molecular-weight core subunit PsbM. In *Euglena*, PsbM contains an extended C-terminal stromal motif that is absent from its counterparts in green algae and land plants. This extension enables additional interactions with neighboring core subunits, including PsbA, PsbB, PsbC, PsbL, PsbT and the symmetry-related PsbM at the dimer interface (Extended Data Table 4), thereby creating a more extensive interaction network within the core complex (Fig. 7). Structural comparison further shows that this stromal extension is absent from the corresponding PsbM proteins of green algae and land plants, emphasizing that the expanded interaction network in *Euglena* represents a lineage-specific modification of the PSII dimer interface (Fig. 7). Given the known roles of PsbM and neighboring low-molecular-weight subunits in PSII stability and acceptor-side function^33–38^, this lineage-specific extension is likely to reinforce the local architecture of the core and may influence its functional properties. Consistent with this interpretation, previous work showed that loss of PsbM perturbs the acceptor-side region of PSII^38^, suggesting that the expanded PsbM interaction network in *Euglena* may indirectly affect the geometry of the plastoquinone-binding environment at the Q_A_ and Q_B_ sites.

**Fig. 7.**
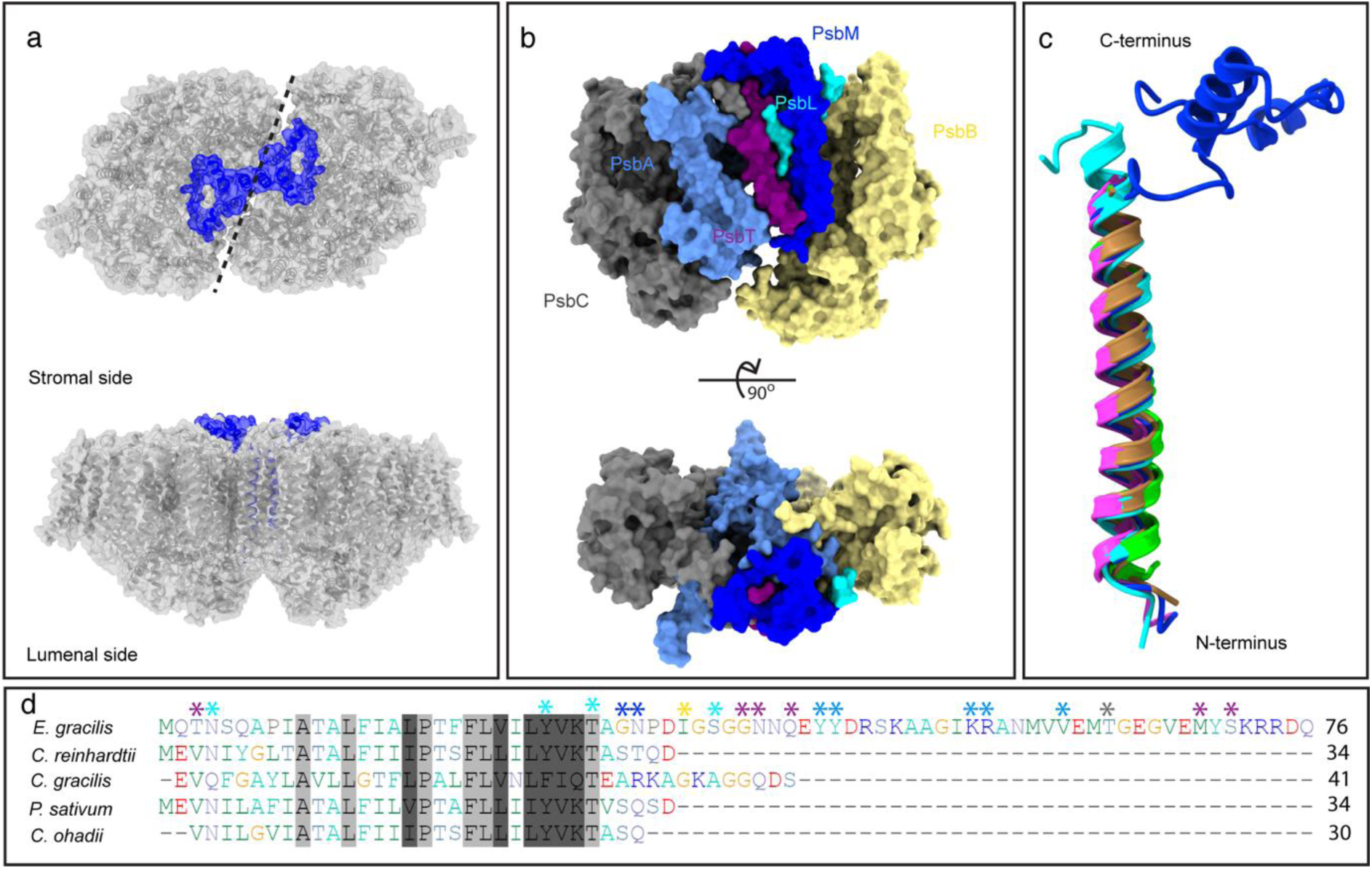
Structure and interactions of PsbM protein in *E. gracilis*. **a** Localization of transmembrane and stromal domains of PsbM protein within the core complex. The C-terminal stromal motif of *E. gracilis* PsbM protein extends over large surface of core complex. **b** Association of PsbM protein with core subunits PsbA, B, C, L, M, T. **c** Structural alignment of *E. gracilis* PsbM protein (blue) with its counterparts in *C. reinhardtii* (6kac, green), *C. gracilis* (6jlu, cyan), *P. sativum* (5xnl, magenta), and *C. ohadii* (9hd7, brown). **d** Alignment of amino acid sequences of PsbM protein from *E. gracilis*, *C. reinhardtii*, *C. gracilis*, *P. sativum*, and *C. ohadii.* The conserved and chemically similar amino acids are depicted in light and dark gray, respectively, whereas the substituted residues are colored. A longer C-terminal motif can be visualized in the amino acid sequence comparison. The PsbM residues involved in the interactions with the core proteins are marked with asterisks with colors corresponding to the core subunits in (**b**).

Additional differences are also apparent in the peripheral antenna system. The trimeric LHCII complexes contain extra chlorophyll-binding sites relative to canonical green algal PSII, including additional Chl *a*715 sites in the S-and L-trimers, similar to those observed in our recent PSII structure from *C. ohadii*^11,12^, whereas the corresponding sites are absent from M-LHCII (Extended Data Figs. 12 and 13). Sequence comparisons further reveal variation in the C-terminal chlorophyll-binding motifs of the trimeric antenna chains (Extended Data Fig. 14), and structural comparison indicates that local differences in C-terminal loop architecture can influence occupancy of these sites (Extended Data Fig. 13). In particular, the longer C-terminal loop in M-LHCII is not compatible with Chl *a*715 binding. These changes indicate that the pigment network of the *Euglena* antenna has been modified beyond the replacement of Lhcb5 by LhcE9. In parallel, structural comparison with algal PSII revealed local positional shifts of Lhcb4 and the M- and L-LHCII trimers, indicating that the geometry of the outer antenna layer has also been adjusted (Extended Data Fig. 15). Although these modifications do not alter the overall architecture of the supercomplex, they support the view that *Euglena* PSII has undergone multiple local adaptations in antenna organization and connectivity.

A related difference concerns the organization of the monomeric antenna region around Lhcb4. Sequence comparison indicates that *Euglena* Lhcb4 lacks the canonical serine residue corresponding to the STT7-dependent phosphorylation site found in green algal Lhcb4 proteins, including *C. reinhardtii*^39,40^ (Extended Data Fig. 16). Similarly, a recent structural report on the *C. ohadii* PSII supercomplex identified substitution at the equivalent site in Lhcb4 and linked it to the apparent absence of state transitions in that alga^12^. Combined with the stable association of LhcE9 and previous evidence for dynamic LhcE1-4 antenna behavior in *Euglena*^32^, this observation is consistent with the idea that excitation balancing in *Euglena* relies on mechanisms distinct from the canonical state transitions described in Viridiplantae^39,41–43^. In this context, the stable positioning of LhcE9 may provide a plausible docking site for the LhcE1-4 antenna complex proposed previously in *Euglena*^32^, although the present structure does not visualize an additional LhcE antenna bound to PSII.

Taken together, these features show that the *Euglena* PSII supercomplex preserves the general green-lineage PSII scaffold but introduces multiple lineage-specific modifications in both the core and antenna regions. Beyond the replacement of canonical Lhcb5 by LhcE9, the extended PsbM architecture, altered pigment organization and local shifts of the peripheral antenna together indicate that *Euglena* has remodeled the antenna-core system at several levels while maintaining the overall supramolecular framework of PSII.

### Diversification of PSII antenna architecture in a secondary plastid lineage

The *Euglena gracilis* PSII supercomplex retains the general green-lineage architecture of dimeric PSII with associated monomeric and trimeric antennae but introduces a distinct antenna solution through replacement of canonical Lhcb5 by the euglenophyte-specific protein LhcE9. This substitution is not merely compositional. LhcE9 occupies the canonical Lhcb5 position but adopts a markedly different binding orientation and establishes a distinct interaction pattern with the PSII core, particularly with CP43. Thus, *Euglena* preserves the overall PSII scaffold while remodelling a key antenna-core interface.

A central implication of this reorganization is its effect on spectral properties. Long-wavelength chlorophyll states have been associated predominantly with PSI and its peripheral antenna, whereas PSII has generally been viewed as more spectrally constrained^17–26^. Our combined structural, spectroscopic, mutagenesis and proteomic analyses identify LhcE9 as the stably bound PSII antenna subunit most closely associated with the far-red state in the *Euglena* supercomplex. Importantly, the data do not suggest that *Euglena* PSII simply reproduces a canonical PSI red site. Rather, they indicate that a related long-wavelength design principle has been implemented in a distinct PSII antenna context, in which both ligand identity and local chlorophyll geometry appear to shape the spectral outcome. Also, unlike cyanobacterial systems, where far-red light harvesting is achieved through incorporation of alternative chlorophyll species such as chlorophyll *f* or *d* into the PSII core^17,18^, the far-red-associated properties of *Euglena* PSII supercomplex likely emerge from modulation of the protein environment of chlorophyll *a* within the lineage-specific LhcE9 antenna.

The FRET analysis further shows that the unusual architecture of *Euglena* PSII has functional consequences for excitation energy transfer. Although the overall energy-transfer network remains broadly similar to that of green algal PSII^9–12^, two particularly fast routes distinguish the complex: a PsbX-mediated connection from the L-trimer to CP47 and a direct LhcE9-to-CP43 pathway. These features indicate that the lineage-specific antenna composition of *Euglena* PSII is not a passive variation in subunit content but is integrated into the functional coupling between the peripheral antenna and the core. In this context, the strong coupling of LhcE9 to CP43 may be especially significant, as it places this antenna in a privileged position for delivering excitation into the core and potentially for supporting additional LhcE-based antenna associations^32^.

Beyond LhcE9, the *Euglena* PSII structure reveals additional lineage-specific modifications, including an extended PsbM architecture, altered pigment organization and local adjustments in antenna geometry. Collectively, these features indicate that the diversification of *Euglena* PSII is not limited to the replacement of a single antenna subunit but reflects broader remodelling of the antenna-core system. The structure therefore provides a mechanistic framework for understanding how secondary plastid lineages can diversify photosystem organization without altering the conserved photochemical core of oxygenic photosynthesis^1,27,28^. More broadly, the results show that the range of structural solutions available to PSII antenna evolution is greater than previously appreciated and includes the incorporation of long-wavelength light harvesting into a non-canonical PSII antenna module^7,17,18^.

## Methods

### Strain and growth conditions

The *Euglena gracilis* strain Z (ATCC-12716) was used in this study. Cells were initially grown for 3 days in Tris-acetate-phosphate (TAP) medium^44^ supplemented with vitamins (biotin 10^−7^ %, vitamin B12 10^−7^ %, and vitamin B1 2 × 10^−5^ % w/v). Cultures were maintained at 23 °C under continuous white LED illumination (400–700 nm) at a photon flux density of 45 µmol photons m^−2^ s^−1^. Cells were subsequently transferred to Tris-minimal phosphate (TMP) medium (pH 7.0)^44^ supplemented with the same vitamins and adjusted to an initial density of 2 × 10^5^ cells mL^−1^. Cultures were grown for an additional 10 days under identical conditions and harvested during the logarithmic growth phase by centrifugation at 800 × *g* for 10 min.

### CRISPR-Cas9 mutagenesis

Targeted disruption of the *LHCB4* (CP29) gene was performed using CRISPR-Cas9. Two sgRNAs were designed to promote large deletions via non-homologous end joining. Synthetic Alt-R crRNAs (Integrated DNA Technologies) targeting the sequences CATCCGGGGAACATTCATCA and GTTGCTCCTGACTACCTGGA were assembled with Cas9 protein (Alt-R S.p. HiFi Cas9 Nuclease V3, Integrated DNA Technologies) to generate ribonucleoprotein complexes. Mutagenesis was performed following the protocol of Nagamine et al. (2024)^45^ using a mixture of the two ribonucleoprotein complexes. Mutant candidates were screened by genomic PCR using the primer pair CP29-cDNA-21Fw (CAGGTTCACCAAAATGTATTCTGAG) and CP29-cDNA-582Rv (GAGGGTGATGTCAGCAGTTCCAGTG). PCR products of approximately 733 bp corresponded to the wild-type allele, whereas a fragment of ∼342 bp indicated the expected large deletion. Two independent knockout lines (CP29-KO-1 and CP29-KO-4) displaying only the shorter amplification product were selected for further analysis (Extended Data Fig. 17).

### Membrane Preparation

Total membrane fractions were prepared as previously described by Yadav et al. (2017)^46^, with minor modifications. Briefly, cells were disrupted in SHE buffer at pH 7.3, containing 250 mM sucrose, 1 mM EDTA, 1 mM phenylmethylsulfonyl fluoride (PMSF), and 50 μg/mL tosyl-L-lysyl chloromethyl ketone (TLCK), using a microprobe sonicator (Fisherbrand™ FB505, Fisher Scientific). Sonication was performed as three 10-s pulses at 30% amplitude, with 60-s intervals between pulses. Membranes were then isolated by differential centrifugation, with the final pelleting step carried out at 17,000 × *g* for 15 min. Protein concentrations were determined using the Bradford assay^47^, and absorbance was measured at 595 nm with a Lambda 265 UV-VIS spectrophotometer (PerkinElmer).

### Clear-Native Electrophoresis

Membrane proteins were solubilized with n-dodecyl-α-D-maltoside (α-DDM, 2%) at a detergent-to-protein ratio of 4 g g^−1^ in Tris-based solubilization buffer 50 mM Tris-HCl, 1.5 mM MgSO4, 100 mM NaCl, 10% glycerol, 1 mM PMSF), and 50 μg/mL TLCK at pH 8.4. After incubation for 30 min at 4 °C, insoluble material was removed by centrifugation (21,130 × *g*, 20 min). Supernatants were analyzed by high-resolution clear-native PAGE (hrCN-PAGE) on 4-12 % acrylamide gradient gels according to Wittig et al. (2007). Electrophoresis was performed using Bis-Tris/Tricine buffer (5 mM Tricine, 1.5 mM Bis-Tris, pH 7) systems supplemented with low concentrations of sodium deoxycholate (0.05%) and α-DDM (0.02%) to preserve photosynthetic complexes. Following electrophoresis, acrylamide gel pieces corresponding to the largest PSII supercomplex (band B1, see **Fig. 1a**) were excised from the gel and incubated in the solubilization buffer containing 0.01 % α-DDM. Protein complexes were extracted at 4 °C for 24 h under gentle agitation prior to spectroscopic analysis.

### Spectroscopic measurements

Absorbance and fluorescence spectra were recorded from individual excised CN-PAGE bands. Absorbance spectra at room temperature were measured using a BLUE-Wave miniature spectrometer (StellarNet Inc.) equipped with tungsten-halogen and blue light sources. Fluorescence emission spectra were recorded using a USB2000+ spectrometer (Ocean Optics) coupled to a CCD detection unit (Beambio) with excitation at 470 nm. Spectra were acquired both at room temperature and at 77 K. Following data acquisition, excised gel bands were stored at –80 °C.

### Pigment extraction and HPLC analysis

Pigments were extracted from individual excised CN-PAGE bands following the procedure described by Miranda-Astudillo et al. (2025)^32^. Pigment separation and quantification by HPLC were performed according to the method of Berne et al. (2018)^48^.

### Quantitative proteomics and statistical analysis

Protein digestion and LC-MS/MS acquisition were performed at the de Duve Institute’s MASSPROT core facility platform. Protein bands visualized by CN-PAGE were manually excised, in-gel digested with trypsin, and peptides were extracted with 0.1% TFA in 65% ACN before drying in a SpeedVac. Peptides were resuspended in solvent A (0.1% TFA, 2% ACN), loaded onto a reversed-phase pre-column (Acclaim PepMap 100, Thermo Scientific), and separated in backflush mode on an Acclaim PepMap RSLC analytical column (0.075 × 250 mm, Thermo Scientific) using a Vanquish Neo system at 300 nL/min. The gradient was 4-27.5% solvent B (0.1% FA, 98% ACN) over 40 min, 27.5-50% over 20 min, 50-95% over 10 min, followed by 10 min at 95%. Peptides were analyzed on an Orbitrap Fusion Lumos Tribrid mass spectrometer coupled online to the nano-LC. MS1 spectra were acquired in the Orbitrap at 120,000 resolution over m/z 375–1800, with an AGC target of 4 × 10^5^, maximum injection time of 50 ms, and electrospray voltage of 2.1 kV. Data-dependent MS/MS acquisition was performed over 3 s for ions above 2 × 10⁴ counts, using HCD at 30, detection in the linear ion trap, and 40 s dynamic exclusion. MS2 spectra were acquired with an AGC target of 5 × 10^4^ and dynamic maximum injection time. MS/MS data were searched with Sequest HT in Proteome Discoverer 2.5 SP1 against a home-made photosynthetic protein database^32^. Trypsin was specified as the enzyme, allowing up to two missed cleavages, four modifications per peptide, and charge states up to 5. Precursor and fragment mass tolerances were set to 10 ppm and 0.1 Da, respectively. Variable modifications included Met oxidation (+15.995 Da) and N-terminal pyro-Glu formation from Gln (−17.027 Da) or Glu (−18.011 Da). FDR was assessed with Percolator and set to 1% at the protein, peptide, and modification-site levels. Relative protein abundance was estimated by label-free spectral counting. To correct for technical variability, peptide-spectrum match (PSM) counts were normalized using the mean spectral density of each replicate. For each sample, the total number of identified proteins (N) and the sum of all PSMs (P) were determined to calculate the mean spectral density (S = P / N). Individual protein PSM values were then normalized by this factor (PSM_norm = PSM_raw / S) to ensure inter-sample comparability.

Differential protein abundance between CP29 knockout lines and wild-type samples was expressed as log_2_ fold change values calculated from triplicate analyses. Statistical significance was evaluated using a two-tailed Welch’s t-test, with proteins considered significantly differentially abundant when p < 0.05 and |log_2_ fold change| > 1. Data visualization was performed in R using volcano plots with automated label placement implemented through the ggrepel algorithm. Mass spectrometry results are derived from three independent analyses.

### Sample preparation from cryo-EM and data collection

The sample of the C_2_S_2_M_2_L_2_ PSII-LHCII supercomplex was obtained from the native-PAGE bands. Fifty-five gel bands corresponding to PSII-LHCII supercomplex were excised and rapidly frozen in liquid nitrogen. The frozen bands were crushed to fine powder in liquid nitrogen and mixed with elution buffer containing 50 mM Tris-HCl, 1.5 mM MgSO_4_, protease inhibitor (complete EDTA-free, Roche) 0.01% α-DDM, pH 7.2. The PSII supercomplexes were spontaneously eluted from the gel powder overnight with continuous spinning at 4 °C. The gel debris was removed by centrifugation at 30,000 × *g* for 1hr. The supernatant containing the eluted PSII supercomplexes was recovered, and the sample was concentrated using Amicon filter with a 100 kDa cutoff at recommended speed, until a final sample volume of 50 μl and a concentration of 3 mg·ml^-1^.

The concentrated sample was employed for vitrification of the specimen for cryo-EM analysis. A 3.5 μl sample volume was applied on carbon-coated holey support film type R2/1 on 200 mesh copper grids (Quantifoil) which were glow discharged with a PELCO easiGlow, 15 mA, grid negative, at 0.4 mbar and 25 s glowing time. The sample was vitrified at 4°C temperature and 95% humidity with a Vitrobot Mark IV (Thermo Fisher Scientific). The grids were blotted using standard Vitrobot Filter Paper (Grade 595 ash-free filter paper ø55/20 mm) for 4 s and blot force was set to 0. The vitrified grids were clipped and loaded into a Thermo Fisher Scientific Glacios 200 keV transmission electron microscope, equipped with a Falcon 4i direct electron detector. A dataset comprising of 16,497 movies was acquired in linear mode, with a total electron exposure dose of 60 e^-^·Å^-2^ using the EPU (Thermo Fisher Scientific) software at 150 kx magnification and a pixel size of 0.948 Åpx^-1^.

### Processing of cryo-EM data and reconstruction of map

The collected cryo-EM data were imported to cryoSPARC v4.3.1^49^ for single-particle analysis. In the first step, the micrographs were pre-processed for path motion correction and CTF estimation. Following the pre-processing, the PSII particles were picked partially in manual mode and automatically using a blob picker. The picked particles were extracted with a box size of 512 pixels and subjected to 2D classification to remove the non-specific and junk particles. An initial set of refined 24,617 particles was subjected to *ab initio* 3D reconstruction with a C1 symmetry. The *ab initio* 3D map was utilized to create a set of 50 templates that were employed in template-based particles picking mode, providing a heterogenous large set of 6,815,866 particles. The picked particles were iteratively refined by 2D classification until a clean set of 661,916 particles covering all PSII supercomplex views was obtained. Following the 2D classification the obtained set of particles was subjected to 3D refinement with C2 symmetry, resulting in a higher-resolution map at 3.79 Å resolution (FSC = 0.143). To enrich the dataset with homogenous and high-resolution particles, a 3D classification was performed with 4 classes. A total of 354,127 particles were obtained from the initial 3D classification run which were subjected to iterative 2D and 3D classification until a refined set of 267,352 particles was obtained. The particles were subjected to reference motion correction and local refinement taking into consideration their individual CTF parameters, which resulted a final 3D reconstruction at 3.09 Å (FSC = 0.143).

### Model building, refinement and model analysis

Initial fitting of the C_2_S_2_M_2_L_2_ into the cryo-EM map was carried out by rigid-body real-space refinement in Chimera, using the crystal structures of PSII from *Ch. ohadii and Cr. reinhardtii* (PDBs 9HD7 and 6KAD) as starting templates. Local fitting of all chains and ligands into the cryo-EM map was performed in Coot. Amino acids were updated to the protein sequences available at the Uniprot database, three chains namely M, O and Z are unmapped, and only based on available transcriptome data^32^. Starting model for the LhcE9 subunit was constructed in Alphafold 3, while the PsbM subunit was created manually. Finally, 4 diatoxanthin (ET4) and 48 diadinoxanthin (DD6) molecules were placed and refined into carotenoid densities across the whole supercomplex. The resulting model was refined against the 3.1 Å cryo-EM reconstruction (FSC = 0.143) using the Real-Space Refinement module in Phenix, with geometry, secondary-structure, NCS, rotamer, and Ramachandran restraints applied. The validation statistics calculated by MolProbity provided the final score value of 1.83, the overall clash score of 6.45 and zero Ramachandran outliers (Supplementary Table 1). FRET analysis was performed according to Sheng et al. (2019) and Croce and Amerongen (2020).

## Data availability

The cryo-EM map of *E. gracilis* PSII-LHCII supercomplex has been deposited in the Electron Microscopy Data Bank with accession codes EMD-58395. The structure model of C_2_S_2_M_2_L_2_ supercomplex is deposited in the RCSB under the PDB accession code 31GH. Mass spectrometry proteomics data has been deposited to the ProteomeXchange Consortium via PRIDE partner repository with accession code PXD080050.

## Acknowledgements

This work was supported by the Programme Jan Amos Komenský Operational Marie Skłodowska-Curie Actions Czechia (OP JAK MSCA CZ) fellowship to R.A. (project no. CZ.02.01.01/00/22_010/0006945), by the Johannes Amos Comenius Programme – Excellent Research to D.K. and R.K. (project no. CZ.02.01.01/00/22_008/0004624), by the Grant Agency of the Czech Republic (project no. 26-21131S) to D.K. and R.K., by the Belgian Fonds de la Recherche Scientifique (FNRS) to P.C. (grants nos R.M011.26, and J.0025.24 to PC), and by JSPS KAKENHI to M.N. (grant no. JP24K01897) and by PAPIIT-UNAM (Grant: IN217826 to HMA) and IIBO-Institutional Program “Production of biomolecules of biomedical interest in microorganisms” to HMA.. P.C and H.F thank Daivy Nguyen for his valuable technical support, and Didier Vertommen for his contribution to the acquisition of the mass spectrometry data.

H.F is FRIA grantee and P.C is a Research Director of FNRS. P.L.K. was supported by the European Union through funding of the Horizon Europe projects ERA Chair “hot4cryo” no. 101086665 and EIC “SWIRFlex-DT” no. 101257652, the Federal Ministry of Education and Research (Bundesministerium für Bildung und Forschung, Zentrum für Innovationskompetenz program) (grant nos. 03Z22HN23, 03Z22HI2, and 03COV04), the European Regional Development Funds (Europäischer Fonds für regionale Entwicklung) for Saxony-Anhalt (grant nos. ZS/2016/04/78115 and ZS/2024/05/187255), the Deutsche Forschungsgemeinschaft (project nos. 391498659, RTG 2467 and 514901783, SFB 1664 [A04, C04, and D01]), and the Martin Luther University Halle-Wittenberg.

## Authors contributions

R.A., H.M.A., P.C., R.K. designed the study. H.F. separation of pigment-protein complexes. R.A., F.H., H.M.A. sample preparation for cryo-electron microscopy. R.A., F.H. collected the high-resolution cryo-EM data. R.A., P.K. image analysis of cryo-EM data. D.K., R.A., R.K. model building and analysis. H.F. mass spectrometry, spectroscopy, and pigment analyses. M.N., H.F. preparation and analysis of the koLHCB4 mutant. R.K. performed the FRET analysis. R.A., H.F., D.K., P.C., R.K. data interpretation. P.L.K. supervised the cryo-EM work and secured access to the instrumentation. R.A. wrote the major part of the initial draft of the manuscript. H.F., P.C., and R.K. contributed to the drafting and development of the manuscript and to figure preparation. All authors revised and approved the final version.

## Competing interests

The authors declare no competing interests.

## Notes

### Competing Interest Statement

The authors have declared no competing interest.

